# The helminth glycoprotein omega-1 improves metabolic homeostasis in obese mice through type-2 immunity-independent inhibition of food intake

**DOI:** 10.1101/2020.07.03.186254

**Authors:** Hendrik J.P. van der Zande, Michael A. Gonzalez, Karin de Ruiter, Ruud Wilbers, Noemi Garcia-Tardón, Mariska van Huizen, Kim van Noort, Leonard R. Pelgrom, Joost M. Lambooij, Anna Zawistowska-Deniziak, Frank Otto, Arifa Ozir-Fazalalikhan, Danny van Willigen, Mick Welling, Jordan Poles, Fijs van Leeuwen, Cornelis H. Hokke, Arjen Schots, Maria Yazdanbakhsh, P’ng Loke, Bruno Guigas

## Abstract

Type 2 immunity plays an essential role in the maintenance of metabolic homeostasis and its disruption during obesity promotes meta-inflammation and insulin resistance. Infection with the helminth parasite *Schistosoma mansoni* and treatment with its soluble egg antigens (SEA) can induce a type 2 immune response in metabolic organs and improve insulin sensitivity and glucose tolerance in obese mice, yet a causal relationship remains unproven. Here, we investigated the effects and underlying mechanisms of the T2 ribonuclease omega-1 (ω1), one of the major *S. mansoni* immunomodulatory glycoproteins, on metabolic homeostasis. Male C57Bl6/J mice were fed a high-fat diet for 12 weeks followed by bi-weekly injection of SEA, ω1 or vehicle for 4 additional weeks. Whole-body metabolic homeostasis and energy expenditure were assessed by glucose/insulin tolerance tests and indirect calorimetry, respectively. Tissue-specific immune cell phenotypes were determined by flow cytometry. We show that treatment of obese mice with plant-produced recombinant ω1, harboring similar glycan motifs as present on the native molecule, decreased body fat mass and improved systemic insulin sensitivity and glucose tolerance in a time-and dose-dependent manner. This effect was associated with an increase in white adipose tissue (WAT) type 2 T helper cells, eosinophils and alternatively-activated macrophages, without affecting type 2 innate lymphoid cells. In contrast to SEA, the metabolic effects of ω1 were still observed in obese STAT6-deficient mice with impaired type 2 immunity, indicating that its metabolic effects are independent of the type 2 immune response. Instead, we found that ω1 inhibited food intake, without affecting locomotor activity, WAT thermogenic capacity or whole-body energy expenditure, an effect also occurring in leptin receptor-deficient obese and hyperphagic *db/db* mice. Altogether, we demonstrate that while the helminth glycoprotein ω1 can induce type 2 immunity, it improves whole-body metabolic homeostasis in obese mice by inhibiting food intake via a STAT6-independent mechanism.

**Author summary:** The obesity-induced chronic low-grade inflammation, notably in adipose tissue, contributes to insulin resistance and increased risk of type 2 diabetes. We have previously shown that infection with parasitic helminth worms was associated with protection against obesity-related metabolic dysfunctions both in mice and humans. We have also reported that treatment of obese mice with an extract of *Schistosoma mansoni* eggs (SEA) improves insulin sensitivity and glucose tolerance, a beneficial effect that was associated with a helminth-specific type 2 immune response in metabolic organs. Here, we studied the effects of omega-1 (ω1), a single immunomodulatory molecule from SEA, on metabolic health in obese mice, and investigated the role of the host immune response elicited. We found that ω1 induced a helminth-characteristic type 2 immune response in adipose tissue and improved both insulin sensitivity and glucose tolerance in obese mice. Yet, in contrast to SEA, ω1’s immunomodulatory properties were dispensable for its metabolic effects. Instead, we show that ω1 inhibited food intake, a feature accounting for most of the improvements in metabolic health. Together, our findings indicate that helminth molecules may improve metabolic health through multiple distinct mechanisms, and further characterization of such molecules could lead to new therapeutic strategies to combat obesity.

## Introduction

Obesity is associated with chronic low-grade inflammation in metabolic organs (1). This so-called meta-inflammation plays a prominent role in the etiology of insulin resistance and type 2 diabetes (1-3), and is associated with increased numbers of pro-inflammatory macrophages, notably in white adipose tissue (WAT) (4) and liver (5). In WAT, these macrophages mainly originate from newly-recruited blood monocytes that differentiate into pro-inflammatory macrophages upon entering the inflammatory milieu (4) and/or being activated by elevated local concentration of free fatty acids (6). These pro-inflammatory macrophages produce cytokines, such as tumor necrosis factor (TNF) and interleukin 1-beta (IL-1β), which directly inhibit canonical insulin signaling [as reviewed in (2)] and contribute to tissue-specific insulin resistance and whole-body metabolic dysfunctions. In the liver, activation of Kupffer cells, the tissue-resident macrophages, promote the recruitment of pro-inflammatory monocytes and neutrophils which trigger hepatic inflammation and insulin resistance through the production of pro-inflammatory cytokines and elastase, respectively (5, 7, 7). In contrast, a type 2 cytokine environment predominates in lean metabolic tissues under homeostatic conditions, notably in WAT where IL-4, IL-5 and IL-13 produced by type 2 innate lymphoid cells (ILC2s), T helper 2 (Th2) cells and/or eosinophils promote alternatively activated macrophages (AAM) (9, 10). According to the current paradigm, AAMs are the final effector cells of this type 2 immune response, contributing to the maintenance of WAT insulin sensitivity by underlying molecular mechanism(s) that are still largely unknown (2, 11).

Parasitic helminths are the strongest natural inducers of type 2 immunity (12). Interestingly, several studies have reported an association between helminth-induced type 2 immunity and improved whole-body metabolic homeostasis in both humans and rodents [as reviewed in (11)]. We also showed that chronic treatment with *S. mansoni* soluble egg antigens (SEA) promoted eosinophilia, Th2 cells, type 2 cytokines expression and AAMs in WAT, and improved both tissue-specific and systemic insulin sensitivity in obese mice (13). SEA drives dendritic cell (DC)-mediated Th2 skewing at least partly through glycosylated molecules [(14), and reviewed in (15)], particularly the T2 RNase glycoprotein omega-1 [ω1; (16, 17)]. Interestingly, acute treatment with human embryonic kidney 293 (HEK-293)-produced recombinant ω1 was recently shown to decrease body weight and improve whole-body glucose tolerance in obese mice, through ILC2-mediated type 2 immunity and induction of WAT beiging (18). In this study, the metabolic effect of ω1 was reported to be glycan-dependent, yet we have previously shown that the glycosylation pattern of HEK-293-produced ω1 differs significantly from the *S. mansoni* native molecule, which notably harbors immunogenic Lewis-X (Le^X^) glycan motifs (17, 19). By exploiting the flexible N-glycosylation machinery of *Nicotiana benthamiana* plants, we successfully produced large amounts of recombinant ω1 glycosylation variants, either carrying Le^X^ motifs on one of its glycan branches or not (20).

In the present study, we investigate the effects and underlying immune-dependent mechanisms of both SEA and two plant-produced ω1 glycovariants on whole-body metabolic homeostasis in obese mice. Remarkably, we demonstrate that while SEA improved metabolic homeostasis in obese mice through a STAT6-dependent type 2 immune response, recombinant pLe^X^-ω1 did so independent of its type 2 immunity-inducing capacity, by reducing food intake in a leptin receptor-independent manner.

## Methods

### Animals, diet and treatment

All mouse experiments were performed in accordance with the Guide for the Care and Use of Laboratory Animals of the Institute for Laboratory Animal Research and have received approval from the university Ethical Review Board (Leiden University Medical Center, Leiden, The Netherlands; DEC12199) or the Institutional Animal Care and Use Committee (IACUC, New York University School of Medicine, New York, USA; protocol ID IA16-00864). All mice were housed in a temperature-controlled room with a 12 hour light-dark cycle with *ad libitum* access to food and tap water. Group randomization was systematically performed before the start of each experiment, based on body weight, fat mass, and fasting plasma glucose levels. At the end of the experiment, mice were sacrificed through an overdose of ketamine/xylazine.

8-10 weeks old male wild-type (WT) and 7 weeks old male *db/db* mice, both on C57BL/6J background, were purchased from Envigo (Horst, The Netherlands) and housed at Leiden University Medical Center. WT mice were fed a low-fat diet (LFD, 10% energy derived from fat, D12450B, Research Diets, Wijk bij Duurstede, The Netherlands) or a high-fat diet (HFD, 45% energy derived from fat, D12451) for 12 weeks, and d*b/db* mice were fed a chow diet (RM3 (P), Special Diet Services, Witham, UK) throughout the experimental period. SEA was prepared as described previously (21). Recombinant ω1 was produced in *N. benthamiana* plants through transient expression of ω1 alone (pWT-ω1) or ω1 in combination with exogenous glycosyltransferases to yield Le^X^ glycan motifs (pLe^X^-ω1), as described previously (20). SEA, pWT/pLe^X^-ω1 (10-50 μg) or vehicle control (sterile-filtered PBS) were injected i.p. every 3 days for 1 or 4 weeks, as indicated in the legends of the figures. For fast-refeeding experiments, WT HFD-fed mice received an i.p. injection of 50 μg pLe^X^-ω1 or vehicle control after an overnight fast (5pm-9am), followed by refeeding and frequent measurements of food intake and body weight during 24 hours. For assessing the contribution of reduced food intake on the immunometabolic effects of pLe^X^-ω1, WT HFD-fed mice were single-housed and, in a pair-fed group of PBS-injected mice, daily food availability was adjusted to the calorie intake of the pLe^X^-ω1-treated group.

8-10 weeks-old male wild-type (WT) and *Stat6*^-/-^ mice, both on C57BL/6J background, were purchased The Jackson Laboratory (Bar Harbor, ME, USA), housed at New York University School of Medicine, and either put on a HFD (60% energy derived from fat; D12492; Research Diets, New Brunswick, NJ, USA) or LFD (10% energy derived from fat; D12450J; Research Diets) for 10 weeks. To exclude effects of genotype-dependent microbiota differences on metabolic and immunological outcomes, the beddings of WT and *Stat6*^-/-^ mice were frequently mixed within similar diet groups throughout the run-in period. After 10 weeks, HFD-fed mice were randomized as described above and treated every 3 days for 4 weeks with 50 μg SEA, pLe^X^-ω1 or vehicle-control.

### Body composition and indirect calorimetry

Body composition was measured by MRI using an EchoMRI (Echo Medical Systems, Houston, TX, USA). Groups of 4-8 mice with free access to food and water were subjected to individual indirect calorimetric measurements during the initiation of the treatment with recombinant ω1 for a period of 7 consecutive days using a Comprehensive Laboratory Animal Monitoring System (Columbus Instruments, Columbus, OH, USA). Before the start of the measurements, single-housed animals were acclimated to the cages for a period of 48 hours. Feeding behavior was assessed by real-time food intake. Oxygen consumption and carbon dioxide production were measured at 15 minute intervals. Energy expenditure (EE) was calculated and normalized for lean body mass (LBM), as previously described (13). Spontaneous locomotor activity was determined by the measurement of beam breaks.

At sacrifice, visceral white adipose tissue (epidydimal; eWAT), subcutaneous white adipose tissue (inguinal; iWAT), supraclavicular brown adipose tissue (BAT) and liver were weighed and collected for further processing and analyses.

### Isolation of stromal vascular fraction from adipose tissue

eWAT was collected at sacrifice after a 1 minute perfusion with PBS through the heart left ventricle and digested as described previously (13). In short, collected tissues were minced and incubated for 1 hour at 37°C in an agitated incubator (60 rpm) in HEPES buffer (pH 7.4) containing 0.5 g/L collagenase type I from *Clostridium histolyticum* (Sigma-Aldrich, Zwijndrecht, The Netherlands) and 2% (w/v) dialyzed bovine serum albumin (BSA, fraction V; Sigma-Aldrich). The disaggregated adipose tissue was passed through a 100 μm cell strainer that was washed with PBS supplemented with 2.5 mM EDTA and 5% FCS. After centrifugation (350 x g, 10 minutes at room temperature), the supernatant was discarded and the pellet was treated with erythrocyte lysis buffer (0.15 M NH_4_Cl; 1 mM KHCO_3_; 0.1 mM Na_2_EDTA). Cells were next washed with PBS/EDTA/FCS, and counted manually.

### Isolation of leukocytes from liver tissue

Livers were collected and digested as described previously (13). In short, livers were minced and incubated for 45 minutes at 37°C in RPMI 1640 + Glutamax (Life Technologies, Bleiswijk, The Netherlands) containing 1 mg/mL collagenase type IV from *Clostridium histolyticum*, 2000 U/mL DNase (both Sigma-Aldrich) and 1 mM CaCl_2_. The digested liver tissues were passed through a 100 μm cell strainer that was washed with PBS/EDTA/FCS. Following centrifugation (530 x g, 10 minutes at 4°C), the supernatant was discarded, after which the pellet was resuspended in PBS/EDTA/FCS and centrifuged at 50 x g to pellet hepatocytes (3 minutes at 4°C). Next, supernatants were collected and pelleted (530 x g, 10 minutes at 4°C). The cell pellet was first treated with erythrocyte lysis buffer and next washed with PBS/EDTA/FCS. CD45^+^ leukocytes were isolated using LS columns and CD45 MicroBeads (35 μL beads per liver, Miltenyi Biotec) according to manufacturer’s protocol and counted manually.

### Processing of isolated immune cells for flow cytometry

For analysis of macrophage and lymphocyte subsets, both WAT stromal vascular cells and liver leukocytes were stained with the live/dead marker Aqua (Invitrogen, Bleiswijk, The Netherlands) or Zombie-UV (Biolegend, San Diego, CA, USA), fixed with either 1.9% formaldehyde (Sigma-Aldrich) or the eBioscience™ FOXP3/Transcription Factor Staining Buffer Set (Invitrogen), and stored in FACS buffer (PBS, 0.02% sodium azide, 0.5% FCS) at 4°C in the dark until subsequent analysis. For analysis of cytokine production, isolated cells were cultured for 4 hours in culture medium in the presence of 100 ng/mL phorbol myristate acetate (PMA), 1 μg/mL ionomycin and 10 μg/mL Brefeldin A (all from Sigma-Aldrich). After 4 hours, cells were washed with PBS, stained with Aqua, and fixed as described above.

### Flow cytometry

For analysis of CD4 T cells and innate lymphoid cell (ILC) subsets, SVF cells were stained with antibodies against B220 (RA3-6B2), CD11b (M1/70), CD3 (17A2), CD4 (GK1.5), NK1.1 (PK136) and Thy1.2 (53-2.1; eBioscience), CD11c (HL3) and GR-1 (RB6-8C5; both BD Biosciences, San Jose, CA, USA), and CD45.2 (104; eBioscience, Biolegend or Tonbo Biosciences, San Diego, CA, USA). CD4 T cells were identified as CD45^+^ Thy1.2^+^ Lineage^+^ CD4^+^, and ILCs as CD45^+^ Thy1.2^+^ Lineage^-^ CD4^-^ cells, in which the lineage cocktail included antibodies against CD11b, CD11c, B220, GR-1, NK1.1 and CD3.

CD4 T cell subsets and cytokine production by ILCs were analyzed following permeabilization with either 0.5% saponin (Sigma-Aldrich) or eBioscience™ FOXP3/Transcription Factor Staining Buffer Set. Subsets were identified using antibodies against CD11b, CD11c, GR-1, B220, NK1.1, CD3, CD45.2, CD4, Thy1.2, IL-4 (11B11), IL-13 (eBio13A), Foxp3 (FJK-16s; all eBioscience), IL-5 (TRFK5) and IFN-γ (XMG1.2; both Biolegend).

For analysis of macrophages, eosinophils, monocytes and neutrophils, cells were permeabilized as described above. Cells were then incubated with an antibody against YM1 conjugated to biotin (polyclonal; R&D Systems, Minneapolis, MN, USA), washed, and stained with streptavidin-PerCP (BD Biosciences) or streptavidin-PerCP-Cy5.5 (Biolegend), and antibodies directed against CD45 (30-F11, Biolegend), CD45.2, CD11b, CD11c [HL3 (BD Biosciences) or N418 (Biolegend)], F4/80 (BM8; Invitrogen or Biolegend), Siglec-F (E50-2440; BD Biosciences), and Ly6C (HK1.4; Biolegend).

All cells were stained and measured within 4 days post fixation. Flow cytometry was performed using a FACSCanto or LSR-II (both BD Biosciences), and gates were set according to Fluorescence Minus One (FMO) controls. Representative gating schemes are shown in Figure S1 and all antibodies used are listed in Table S1.

### Plasma analysis

Blood samples were collected from the tail tip of 4h-fasted mice (food removed at 9 am) using chilled paraoxon-coated capillaries. Fasting blood glucose level was determined using a Glucometer (Accu-Check; Roche Diagnostics, Almere, The Netherlands) and plasma insulin level was measured using a commercial kit according to the instructions of the manufacturer (Chrystal Chem, Zaandam, The Netherlands). The homeostatic model assessment of insulin resistance (HOMA-IR) adapted to mice (22) was calculated as ([glucose (mg/dl)*0.055] × [insulin (ng/ml) × 172.1])/3857, and used as a surrogate measure of whole-body insulin resistance.

### Glucose and insulin tolerance tests

Whole-body glucose tolerance test (ipGTT) was performed at week 3 of treatment in 6h-fasted mice, as previously reported (13). In short, after an initial blood collection by tail bleeding (t = 0), a glucose load (2 g/kg total body weight of D-Glucose [Sigma-Aldrich]) was administered i.p., and blood glucose was measured at 20, 40, 60, and 90 min after glucose administration using a Glucometer. For *db/db* mice, blood samples were collected at 0, 20, 40 and 90 min after glucose administration, and plasma glucose levels were measured using the hexokinase method (HUMAN, Wiesbaden, Germany).

Whole-body insulin tolerance test (ipITT) was performed determined at week 1 or week 3 of treatment in 4h-fasted mice, as described previously (13). In short, after an initial blood collection by tail bleeding (t = 0), a bolus of insulin (1 U/kg (lean) body mass [NOVORAPID, Novo Nordisk, Alphen aan den Rijn, Netherlands]) was administered i.p., and blood glucose was measured at 20, 40, 60, and 90 min after insulin administration using a Glucometer.

### Statistical analysis

All data are presented as mean ± standard error of the mean (SEM). Statistical analysis was performed using GraphPad Prism version 8 for Windows (GraphPad Software, La Jolla, CA, USA) with unpaired t-test, or either one-way or two-way analysis of variance (ANOVA) followed by Fisher’s post-hoc test. Differences between groups were considered statistically significant at P < 0.05.

## Results

### *S. mansoni* soluble egg antigens (SEA) improve metabolic homeostasis in obese mice by a STAT6-dependent mechanism

In order to investigate the role of type 2 immunity in the beneficial metabolic effects of SEA, we used mice deficient for STAT6 (*Stat6*^-/-^), a key transcription factor involved in signature type 2 cytokines interleukin (IL)-4/IL-13 signaling and maintenance of Th2 effector functions (23, 24). As previously reported (13), we confirmed that chronic treatment with SEA for 4 weeks increased IL-5 and IL-13-expressing Th2 cells (Fig. 1A-B), eosinophils (Fig. 1C) and YM1^+^ AAMs (Fig. 1D) in WAT from HFD-fed obese WT mice while, as expected, this type 2 immune response was abrogated in *Stat6*^-/-^ mice. SEA slightly reduced body weight (Fig. 1E) and similarly affected body composition (Fig. S1A) in both WT and *Stat6*^-/-^ obese mice, without affecting food intake (Fig. 1F). In line with our previous study, we showed that SEA reduced fasting plasma insulin levels (Fig. S1C) and HOMA-IR (Fig. 1G), and improved whole-body glucose tolerance in WT obese mice. Remarkably, this beneficial metabolic effect was completely abolished in *Stat6*^-/-^ mice (Fig. 1H-J), indicating that SEA improves whole-body metabolic homeostasis in obese mice through STAT6-mediated type 2 immunity.

**Figure 1.**
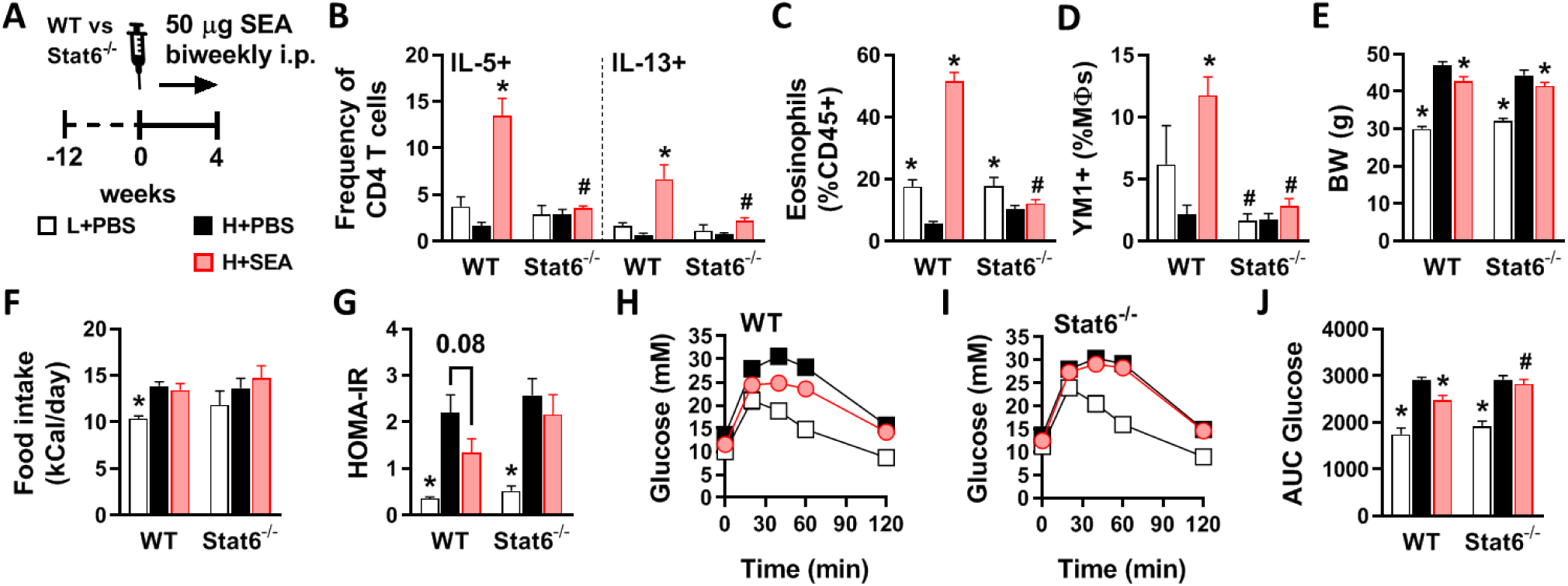
*S. mansoni* soluble egg antigens improve metabolic homeostasis in obese mice by a STAT6-dependent mechanism. WT and *Stat6*-/-mice were fed a LFD (white bars) or a HFD for 12 weeks and next received intraperitoneal injections of PBS (black bars) or 50 μg *S. manson*i soluble egg antigens (SEA; red bars) every 3 days for 4 weeks (*A*). At sacrifice, epididymal WAT was collected and SVF was isolated and analyzed by flow cytometry. The complete gating strategy is shown in Figure S1. Frequencies of IL-5 and IL-13 expressing Th2 cells (*B*) in WAT were determined after PMA/ionomycin/Brefeldin A restimulation. Abundances of eosinophils (*C*) and YM1^+^ macrophages (AAMs; *D*) were determined. Body weight (*E*) was measured after 4 weeks of treatment. Food intake (*F*) was monitored throughout the experimental period. HOMA-IR (*G*) was calculated using fasting blood glucose and plasma insulin levels at week 4. An i.p. glucose tolerance test was performed at week 3. Blood glucose levels were measured at the indicated time points (*H-I*) and the AUC of the glucose excursion curve was calculated (*J*). Data shown are a pool of two independent experiments. Results are expressed as means ± SEM. * *P*<0.05 *vs* HFD, # *P*<0.05 *vs* WT (n = 9-12 mice per group).

### Plant-produced recombinant ω1 glycovariants increase adipose tissue Th2 cells, eosinophils and alternatively-activated macrophages, without affecting innate lymphoid cells

One of the major type 2 immunity-inducing molecules in SEA is the T2 ribonuclease glycoprotein ω1 (17). To study the effect of ω1 on metabolic homeostasis and the role of its immunomodulatory glycans, we generated two recombinant glycosylation variants using glycol-engineered *N. benthamiana* plants: one carrying wild-type plant glycans (pWT-ω1) and one harboring terminal Le^X^ motifs (pLe^X^-ω1; (20)). For both ω1 glycovariants, 4 weeks treatment markedly increased WAT CD4 T cells in HFD-fed obese mice, with pWT-ω1 being more potent than pLe^X^-ω1, while total ILCs were unaffected (Fig. 2A-B). Interestingly, a specific increase in WAT IL-5 and IL-13-expressing Th2 cells was seen for both ω1 glycovariants, while the other CD4 T cell subsets, *i*.*e*. regulatory T cells (Treg) and Th1 cells, were not affected (Fig. 2C). In addition, we confirmed that HFD reduced WAT IL-5^+^/IL-13^+^ ILC2s, as previously reported (9), an effect that was even further pronounced with ω1 glycovariants (Fig. 2D). The type 2 cytokines IL-5 and IL-13 produced by either ILC2s and/or Th2 cells have been reported to maintain WAT eosinophils (9). Congruent with our data on Th2 cells, we found a potent increase in WAT eosinophils upon ω1 treatment that was of similar extent for both glycovariants (Fig. 2E). Finally, both pWT-ω1 and pLe^X^-ω1 increased WAT YM1^+^ AAMs while obesity-associated CD11c^+^ macrophages were not affected (Fig. 2F-G). This ω1-induced WAT type 2 immunity was dose-dependent (Fig. S3) and already observed after one week of treatment, when ILC2s were also not affected (Fig. S4).

**Figure 2.**
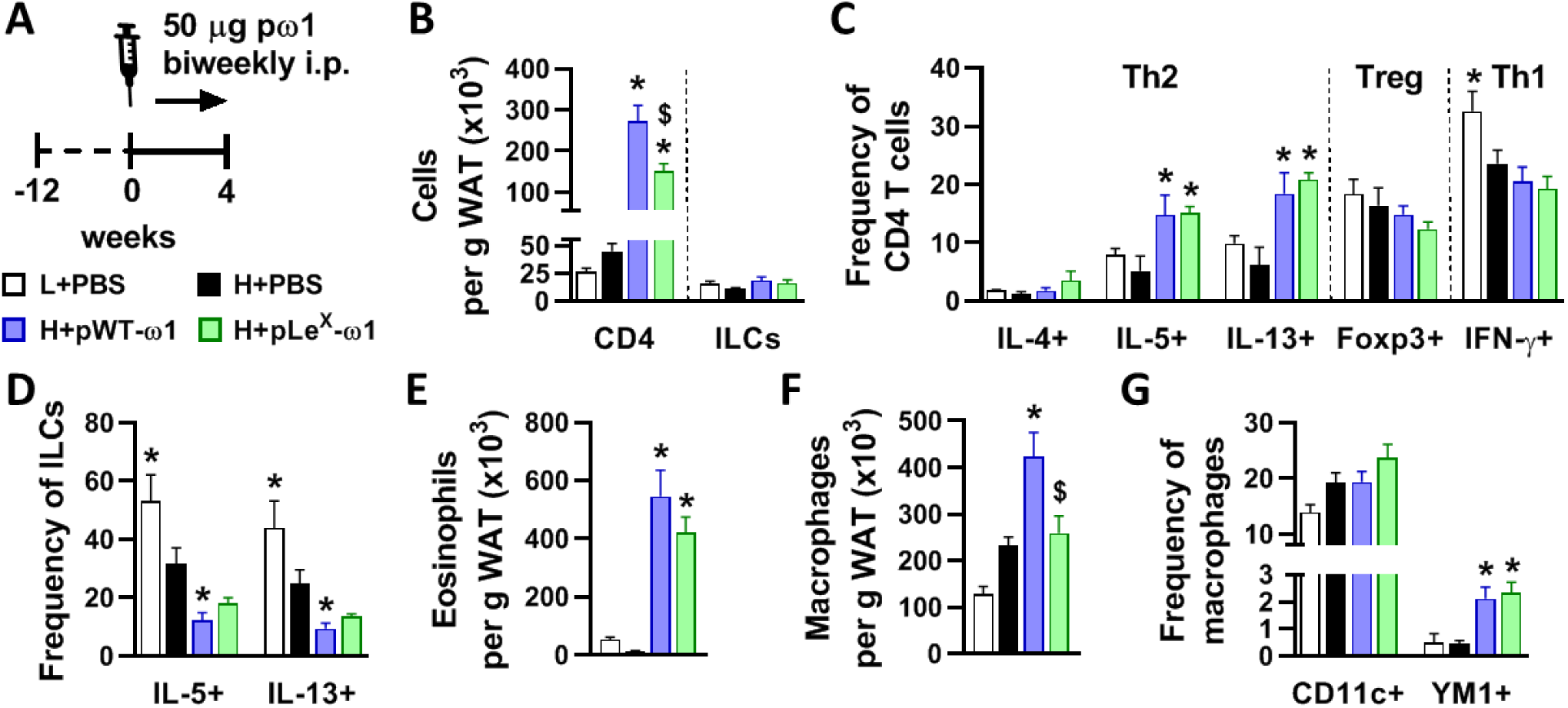
Plant-produced recombinant ω1 glycovariants increase adipose tissue Th2 cells, eosinophils and alternatively-activated macrophages, without affecting innate lymphoid cells. Mice were fed a LFD (white bars) or a HFD for 12 weeks, and next received intraperitoneal injections of PBS (black bars) or either 50 μg pWT-ω1 (blue bars) or 50 μg pLe^X^-ω1 (green bars) every 3 days during 4 weeks (*A*). At the end of the experiment, eWAT was collected, processed and analyzed as described in the legend of Figure 1. Numbers of CD4 T cells, ILCs (*B*), eosinophils (*E*) and macrophages (*F*) per gram tissue were determined. Frequencies of CD4 T helper subsets (*C*) and cytokine-expressing ILCs (*D*) were determined. Percentages of CD11c^+^YM1^-^ and CD11c^-^YM1^+^ macrophages (*G*) were measured. Data shown are a pool of at least two independent experiments. Results are expressed as means ± SEM. * *P*<0.05 *vs* HFD, $ *P*<0.05 *vs* pWT-ω1 (n = 6-19 mice per group in B, E-G, and 3-9 mice per group in C and D).

AAMs are considered the effector cells of WAT type 2 immunity in the maintenance of tissue insulin sensitivity (2), although the mechanisms are not fully understood. Monocyte-derived macrophages can irreversibly be labelled upon tamoxifen administration in *Cx3cr1*^CreERT2-IRES-EYFP^ *Rosa26*^LoxP-stop-LoxP-tdTomato^ (*Cx3cr1*^CreER^ *Rosa26*^tdTomato^) mice, as described elsewhere (25). In order to characterize newly recruited, ω1-induced adipose tissue macrophages (ATMs) during obesity, we performed RNA sequencing on FACS-sorted tdTomato^+^ macrophages from eWAT SVF of obese *Cx3cr1*^CreER^ *Rosa26*^tdTomato^ mice that were treated with PBS or pLe^X^-ω1, the glycovariant that resembles native ω1 most (Fig. S5A). Genes associated with alternative activation, *e*.*g. Tmem26, Slc7a2, Chil3* and *Arg1*, were upregulated (log_2_FC > 2) in ATMs from pLe^X^-ω1-treated mice as compared to controls, while genes associated with pro-inflammatory or obesity-associated macrophages, *e*.*g. Igfbp7, Cxcl12, Bgn, Dcn* and *Cd86*, were downregulated (log_2_FC < −2; Fig. S5B-C). Macrophage function is increasingly recognized to be supported by their metabolism to meet energy demands, and as such, AAMs display increased oxidative phosphorylation (26). In PD-L2^+^ WAT macrophages (Fig. S5D), pLe^X^-ω1 indeed increased mitochondrial mass, while displaying decreased mitochondrial membrane potential and similar total reactive oxygen species production (Fig. S5E-G), a metabolic phenotype in line with alternative macrophage activation.

Similar to WAT, maintenance of insulin sensitivity in the liver is also associated with type 2 immunity (27), whereas obesity-driven activation of Kupffer cells increases the recruitment of pro-inflammatory monocytes and triggers hepatic insulin resistance (2, 7). In our conditions, while ω1 glycovariants increased Th2 cells in the liver, we surprisingly did not find alternative activation of Kupffer cells (Fig. S6A-D). Instead, ω1 glycovariants increased the number of CD11c^+^ pro-inflammatory Kupffer cells, hepatic expression of pro-inflammatory cytokines *Ccl2, Tnf* and *Il1b*, and newly recruited monocytes (Fig. S6D-F), with a more potent effect in pLe^X^-ω1-treated mice. Taken together, these data indicate that both ω1 glycovariants potently induce type 2 immunity in obese mice, triggering an alternative activation profile in WAT, but not liver macrophages.

### ω1 glycovariants reduce body weight, fat mass and food intake, and improve whole-body metabolic homeostasis in obese mice

We next investigated the metabolic effects of ω1 glycovariants and showed that they both induced a rapid and gradual body weight loss in HFD-fed mice (Fig. 3A-B), which was exclusively due to a decrease in fat mass (Fig. 3C). The ω1 glycovariants significantly reduced visceral eWAT mass, but had no or only marginal effects on subcutaneous iWAT, brown adipose tissue (BAT) and liver mass (Fig. S2D). This reduction in fat mass was not due to increased beiging, as ω1 glycovariants neither increased expression of thermogenic genes in both eWAT and iWAT (Fig. S7A-B), nor whole-body energy expenditure (Fig. S7C). In addition, ω1 glycovariants did not affect hepatic steatosis (Fig. S6G-I) but rather increased the expression of fibrotic gene markers (Fig. S6J), without detectable collagen accumulation (Fig. S6K-L). An increase in circulating alanine transaminase levels was also observed (Fig. S6M), indicating that ω1 may also have some cytotoxic effects in the liver, as previously reported (28, 29).

**Figure 3.**
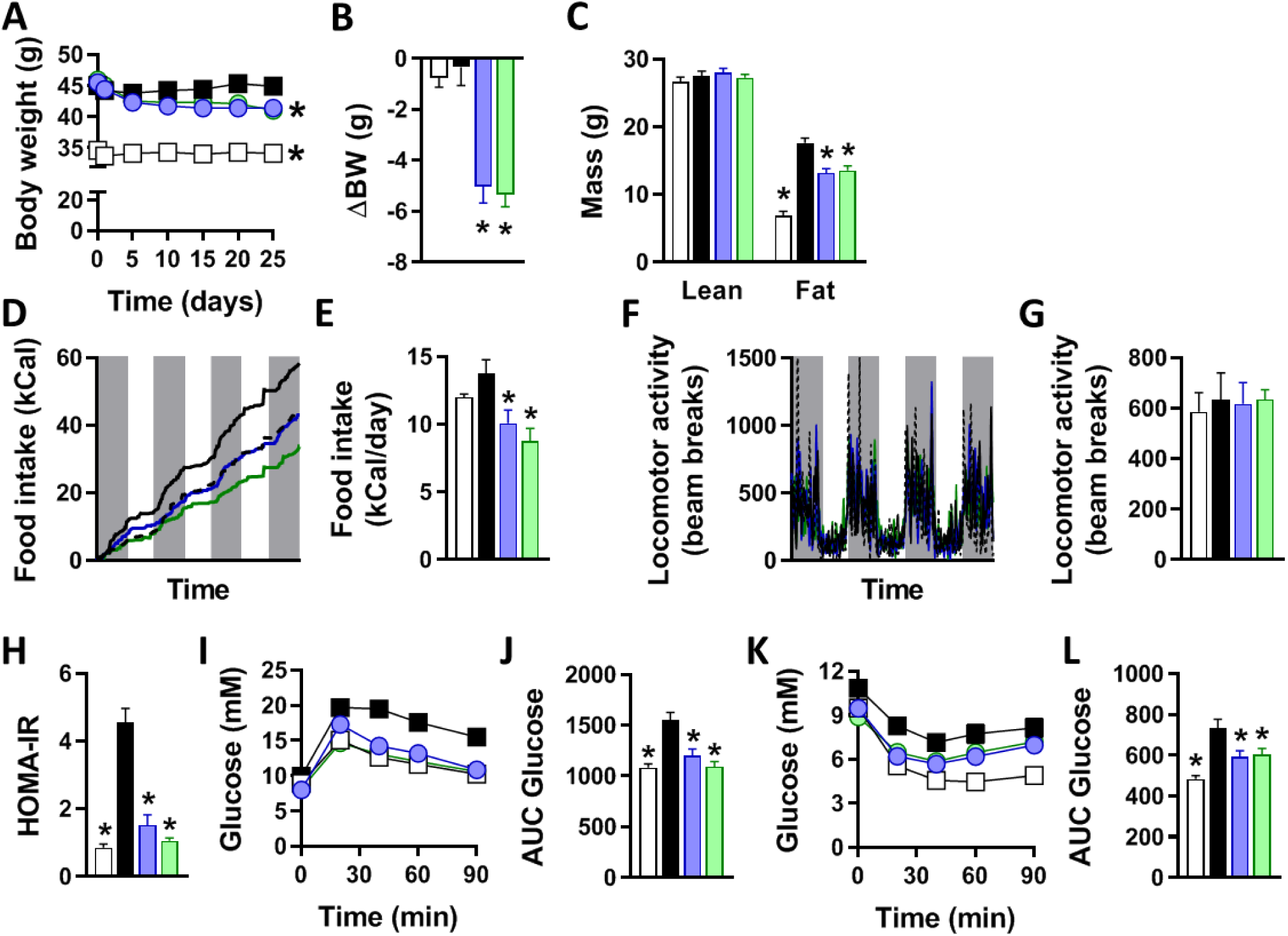
ω1 glycovariants reduce body weight, fat mass and food intake, and improve whole-body metabolic homeostasis in obese mice. Mice were fed a LFD (white bars) or a HFD for 12 weeks, and next received biweekly intraperitoneal injections of PBS (black bars) or 50 μg pWT-ω1 (blue bars) or pLeX-ω1 (green bars) during 4 weeks. Body weight (*A-B*) was monitored throughout the experimental period. Body composition (*C*) was measured after 4 weeks of treatment. Food intake (*D-E*) and locomotor activity (*F-G*) were assessed using fully automated single-housed metabolic cages during the first week of treatment. HOMA-IR at week 4 (*H*) was calculated. Intraperitoneal glucose (*I-J*) and insulin (*K-L*) tolerance tests were performed during week 3. Blood glucose levels were measured at the indicated time points (*I, K*) and the AUC of the glucose excursion curve were calculated (*J, L*). Data shown are a pool of at least 2 independent experiments. Results are expressed as means ± SEM. * *P*<0.05 *vs* HFD (n = 11-20 mice per group in A-C, H-L, and 4-8 mice per group in D-G).

Remarkably, we found that both ω1 glycovariants induced a significant decrease in food intake (Fig. 3D-E), while locomotor activity was not affected (Fig. 3F-G). Treatment with both ω1 glycovariants significantly reduced fasting blood glucose, plasma insulin levels (Fig. S2E-F) and HOMA-IR (Fig. 3H) in obese mice, with a trend towards a stronger effect with pLe^X^-ω1, indicating improved insulin sensitivity. Congruent with these data, we observed a significant improvement in whole-body glucose tolerance (Fig. 3I-J) and insulin sensitivity (Fig. 3K-L) in both pWT-ω1 and pLe^X^-ω1-treated obese mice. Of note, the effects of ω1 glycovariants on food intake, plasma metabolic parameters and whole-body insulin sensitivity were dose-dependent (Fig. S3) and already observed after one week of treatment (Fig. S4). Altogether, these data show that both recombinant ω1 glycovariants improve whole-body metabolic homeostasis in insulin-resistant obese mice.

### pLe^X^-ω1 improves metabolic homeostasis in obese mice by a STAT6-independent mechanism

We next investigated the role of type 2 immunity in the metabolic effects of ω1, using pLe^X^-ω1 as the most potent and native-like glycovariant. As expected, while 4 weeks pLe^X^-ω1 treatment (Fig. 4A) increased WAT Th2 cells, eosinophils and YM1^+^ AAMs in obese WT mice, this type 2 immune response was abrogated in obese *Stat6*^-/-^ mice (Fig. 4B-D). However, treatment with pLe^X^-ω1 still reduced body weight (Fig. 4E-G) and food intake (Fig. 4H), and affected body composition (Fig. S2G) in *Stat6*^-/-^ obese mice to the same extent as in WT mice. In addition, both plasma insulin levels and HOMA-IR were markedly decreased in both genotypes (Fig. S2H-I and Fig. 4I). The improvements in whole-body glucose tolerance (Fig. 4J-L) and insulin sensitivity (Fig. 4M-O) were also still observed in *Stat6*^-/-^ mice, indicating that pLe^X^-ω1’s type 2 immunity-inducing capacity does not play a major role in restoration of metabolic homeostasis in obese mice. Of note, in contrast to its implication in maintenance of WAT metabolic homeostasis, IL-13 signaling has recently also been shown to play a role in the development of liver fibrosis (30, 31). Interestingly, the increase in liver IL-5^+^ and IL-13^+^ Th2 cells in response to pLe^X^-ω1 was also abrogated in *Stat6*^-/-^ mice (Fig. S6N-O), and the expression of fibrotic gene markers were markedly reduced in *Stat6*^-/-^ mice as compared to WT mice (Fig. S6O). Taken together, these results show that pLe^X^-ω1 improves whole-body metabolic homeostasis independent of STAT6-mediated type 2 immunity, while promoting early markers of hepatic fibrosis at least partly through an IL-13-STAT6-mediated mechanism.

**Figure 4.**
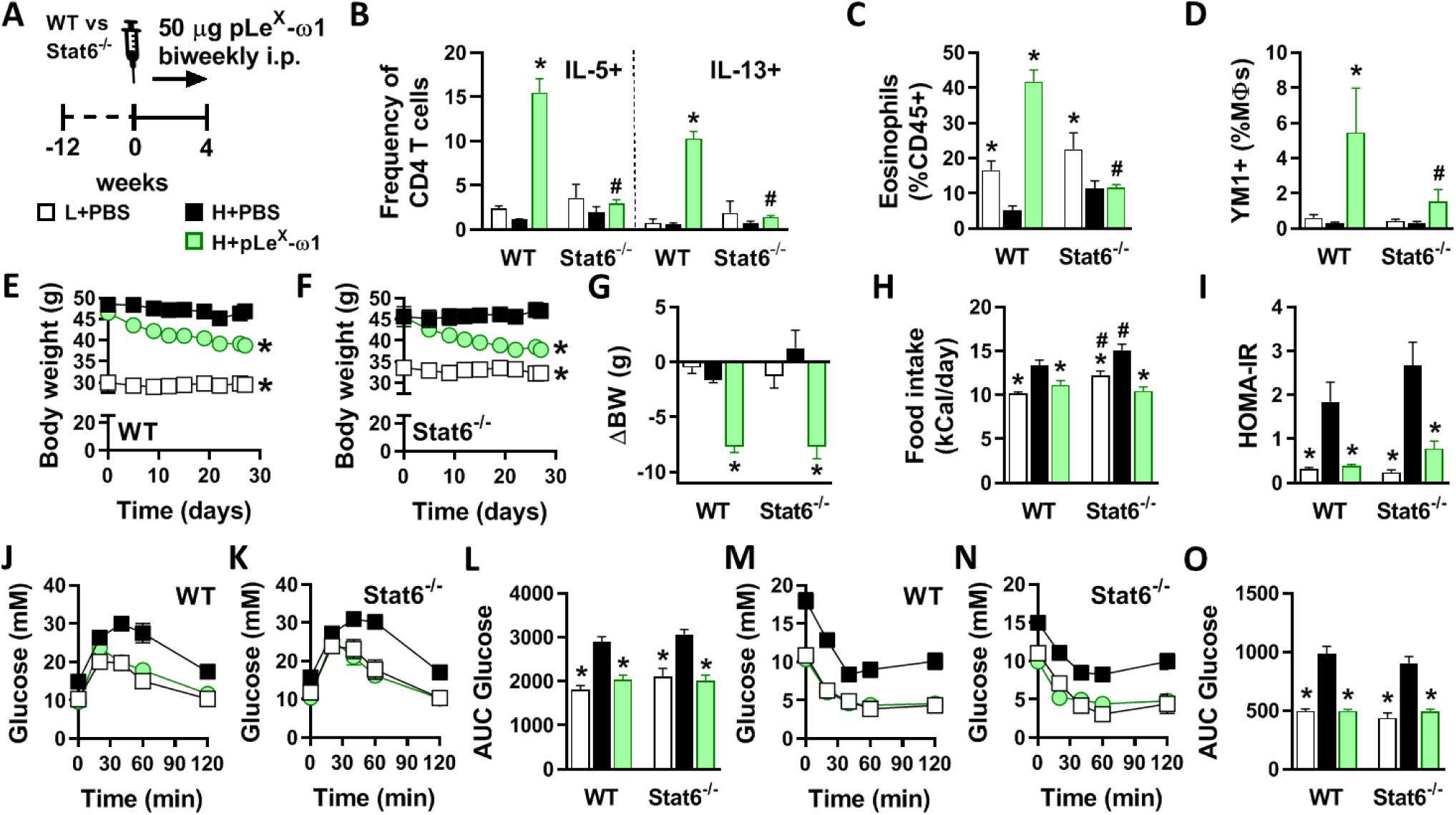
pLe^X^-ω1 improves metabolic homeostasis in obese mice by a STAT6-independent mechanism. WT and *Stat6*-/-mice were fed a LFD (white bars) or a HFD for 12 weeks and next received biweekly intraperitoneal injections of PBS (black bars) or 50 μg pLe^X^-ω1 (green bars) during 4 weeks (*A*). At the end of the experiment, eWAT was collected, processed and analyzed as described in the legend of Figure 1. The frequencies of cytokine-expressing CD4 T cells (*B*) were determined. Abundances of eosinophils (*C*) and YM1^+^ macrophages (*D*) were determined. Body weight (*E-G*) and food intake (*H*) were monitored throughout the experimental period. HOMA-IR at week 4 (*I*) was calculated, and intraperitoneal glucose (*J-L*) and insulin (*M-O*) tolerance tests were performed as described in the legend of Figures 1 and 3. Results are expressed as means ± SEM. \**P*<0.05 *vs* HFD, # *P*<0.05 *vs* WT (n = 3-5 mice per group).

### pLeX-ω1 improves metabolic homeostasis through leptin receptor-independent inhibition of food intake in obese mice

As ω1 significantly reduced food intake in obese mice, we next investigated its impact on feeding behavior. We found that a single intraperitoneal injection of pLe^X^-ω1 in overnight fasted obese mice markedly reduced food intake during refeeding for at least 24 hours, resulting in decreased body weight gain as compared to PBS-injected mice (Fig. 5A-C). To determine whether reduced food intake drives the beneficial metabolic effects of ω1, we treated HFD-fed mice with pLe^X^-ω1 or PBS, and included a pair-fed group that received daily adjusted HFD meals based on the food intake of the pLe^X^-ω1-treated animals (Fig. 5D). While pLe^X^-ω1 expectedly induced IL-13^+^ Th2 cells, eosinophils and YM1^+^ AAMs in WAT, reducing caloric intake in pair-fed mice did not affect WAT type 2 immunity (Fig. 5E-G). Yet, food restriction in the pair-fed group decreased body weight, fasting blood glucose and plasma insulin levels, and HOMA-IR as well as whole-body glucose tolerance to the same extent as in pLe^X^-ω1-treated animals (Fig. S2K-L and Fig. 5H-I).

**Figure 5.**
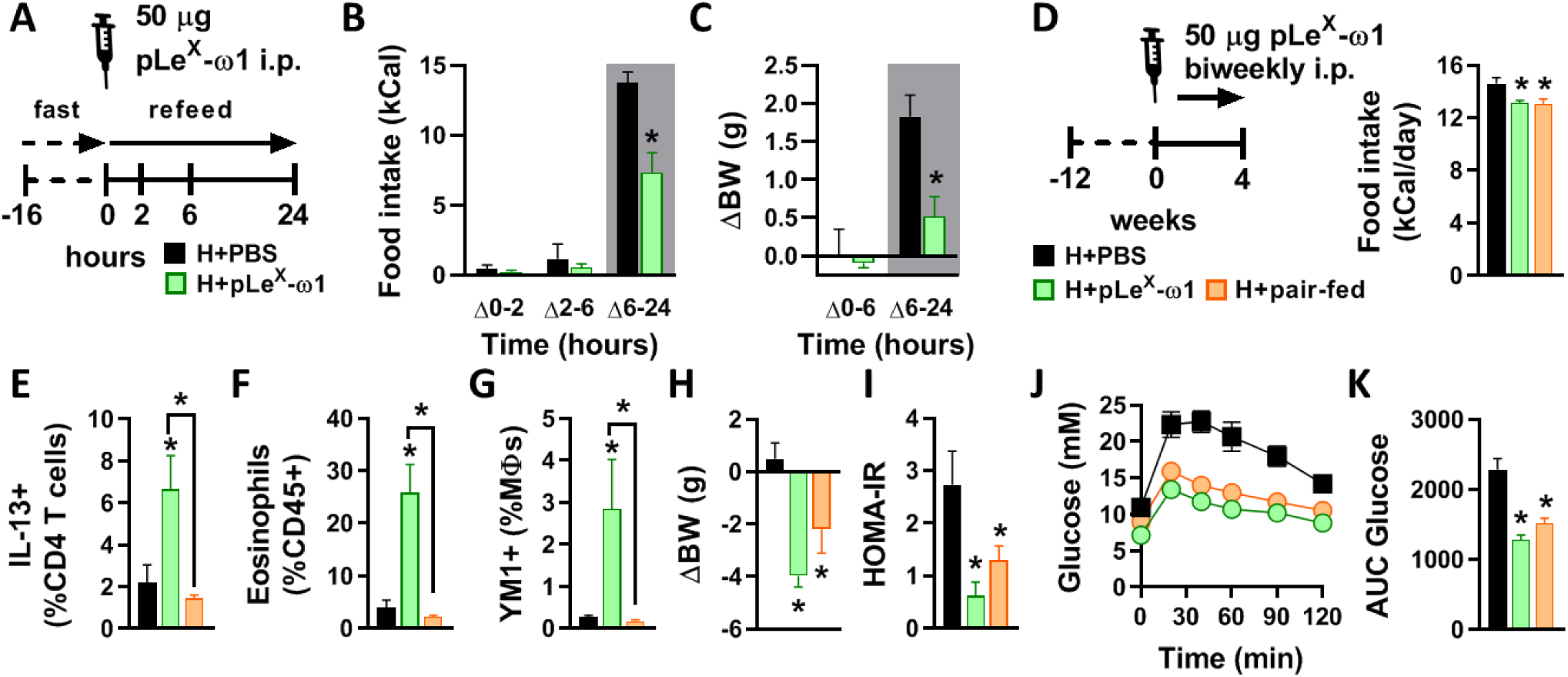
pLe^X^-ω1 inhibits fasting-induced refeeding and improves metabolic homeostasis through inhibition of food intake in obese mice. Mice were fed a HFD for 12 weeks and fasted overnight prior to intraperitoneal injections of either PBS (black bars) or 50 μg pLe^X^-ω1 (green bars; *A*). Food intake (*B*) and body weight changes (*C*) were next monitored during 24 hours after refeeding. Mice were fed a HFD for 12 weeks, single-housed, and next received biweekly intraperitoneal injections of PBS or 50 μg pLe^X^-ω1 during 4 weeks. In one group (H+pair-fed; orange bars), the amount of food available for PBS-treated mice was adjusted daily in order to match the food intake of the pLe^X^-ω1-treated group (*D*). At the end of the experiment, eWAT was collected, processed and analyzed as described in the legend of Figure 1. The frequencies of IL-13-expressing CD4 T cells (*E*), eosinophils (*F*) and YM1^+^ macrophages (*G*) were determined. Body weight change (*H*) was determined after 4 weeks. HOMA-IR (*I*) was calculated at week 4 and an i.p. glucose tolerance test (*J-K*) was performed at week 3, as described in the legend of Figure 1. Results are expressed as means ± SEM. \**P*<0.05 *vs* HFD or as indicated (n = 3-5 mice per group).

The central regulation of feeding behavior and whole-body energy homeostasis involves complex neuronal networks, notably in the hypothalamus and brain stem. (32, 33). To investigate whether pLe^X^-ω1 accumulates in the brain to directly regulate hypothalamic neurons controlling food intake, we performed in vivo imaging experiments with pLe^X^-ω1 conjugated to a hybrid tracer (^111^In-DTPA-Cy5-pLe^X^-ω1). Both Single Photo Emission Computed Tomography (SPECT) imaging and radioactivity biodistribution revealed that ^111^In-DTPA-Cy5-pLe^X^-ω1 mainly accumulated in abdominal organs, *e*.*g*. adipose tissues, liver and intestines, and peritoneal draining lymph nodes, whereas no substantial amounts of radioactivity could be detected in the hypothalamus and other brain regions 24h after tracer administration (Fig. S8A-B). Hence, ω1 does not distribute to the brain and likely regulates food intake through peripheral effects.

The hypothalamus and brain stem also integrate signals from both the enteric nervous system and circulating hormones derived from adipose tissue and other peripheral tissues. Leptin is by far the best studied peripheral hormone that regulates food intake, increasing satiety by triggering STAT3-mediated pathways in the hypothalamic arcuate nucleus (32). In order to study the role of leptin signaling in the metabolic effects of pLe^X^-ω1, we used leptin receptor-deficient *db/db* mice that are hyperphagic and naturally develop obesity and severe metabolic dysfunctions (34). In this model, pLe^X^-ω1 also increased WAT IL-13^+^ Th2 cells, eosinophils and YM1^+^ AAMs (Fig. 6A-D). Furthermore, pLe^X^-ω1 still reduced body weight (Fig. 6E-F), fat mass gain (Fig. S2M-N) and food intake (Fig. 6G), indicating that leptin signaling is not involved in the anorexigenic effect of ω1. Lastly, plasma insulin levels (Fig. S2O-P), HOMA-IR (Fig. 6H) and whole-body glucose tolerance and insulin sensitivity (Fig. 6I-L) were still significantly improved.

**Figure 6.**
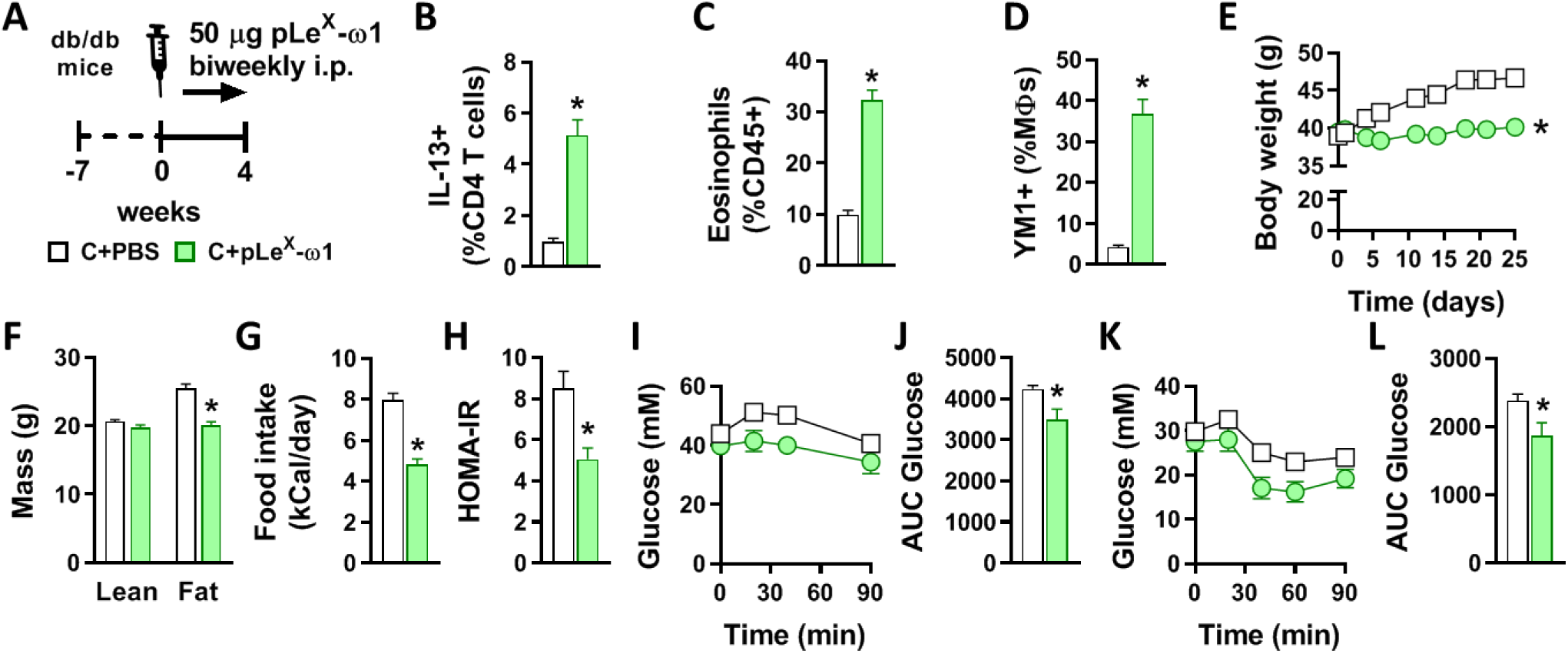
The metabolic effects of pLe^X^-ω1 are independent of leptin signaling in hyperphagic obese mice. 7 weeks-old obese *db/db* mice received biweekly intraperitoneal injections of PBS (white bars) or 50 μg pLe^X^-ω1 (green bars) during 4 weeks (*A*). At the end of the experiment, eWAT was collected, processed and analyzed as described in the legend of Figure 1. The frequencies of IL-13-expressing CD4 T cells (*B*), eosinophils (*C*) and YM1^+^ macrophages (*D*) were determined. Body weight (*E*) and food intake (*G*) were monitored throughout the experimental period, and body composition (*F*) was measured after 4 weeks. Intraperitoneal glucose (*H-I*) and insulin (*J-K*) tolerance tests were performed as described in the legend of Figures 1 and 3. Results are expressed as means ± SEM. \**P*<0.05 *vs* PBS (n = 5-6 mice per group).

Collectively, our results show that ω1 improves whole-body metabolic homeostasis independent of its type 2 immunity-inducing capacity, but by inhibiting food intake through a leptin receptor-independent mechanism.

## Discussion

Obesity-associated metaflammation promotes insulin resistance, while metabolic homeostasis is maintained by type 2 immunity (1). Since helminths are well known for inducing a potent type 2 immune response, their putative beneficial effects on insulin sensitivity and glucose homeostasis, together with the identification of specific helminth-derived molecules capable of driving such type 2 immune responses, have gained increasing attention (11, 15, 15). The assumption has been that induction of type 2 immunity is the main mechanism by which helminths and helminth-derived molecules can improve metabolic homeostasis. The glycoprotein ω1, a T2 ribonuclease which is secreted from *S. mansoni* eggs, is one of the major immunomodulatory components in SEA and has previously been shown to condition DCs to prime Th2 responses, at least partly through its glycan-mediated uptake and intracellular RNase activity (16, 17). Here, we report that two plant-produced recombinant ω1 glycovariants induced a rapid and sustained reduction in body weight and improved whole-body insulin sensitivity and glucose tolerance in obese mice. This improvement was associated with a strong type 2 immune response in WAT, characterized by a significant increase in Th2 cells, eosinophils and AAMs. Contrary to SEA, ω1 still improved metabolic homeostasis in *Stat6*-deficient obese mice, indicating that its type 2 immunity-inducing capacity does not play a major role. Indeed, we find that ω1 regulates energy consumption independent of leptin receptor signaling, which drives most of its metabolic effects. Altogether, these findings indicate that helminth-derived molecules may act through multiple distinct pathways for improving obesity-associated metabolic dysfunctions and further characterization of these molecules may lead to new therapeutic strategies for combating obesity.

A recent study from Hams et al. reported that acute treatment of HFD-fed obese mice with HEK-293-produced recombinant ω1 induced long-lasting weight loss, and improved glucose tolerance by a mechanism involving IL-33 and ILC2-mediated WAT type 2 immunity and adipose tissue beiging (18). In contrast to this report, while we also observed increased IL-33 mRNA expression in eWAT (*data not shown*), we found no increase in WAT ILC2s after either one, or four weeks of treatment with both plant-produced ω1 glycovariants. Moreover, we did not find evidence of WAT beiging in both eWAT and iWAT from obese mice. Lastly, we also found that STAT6-mediated type 2 immunity was dispensable for the metabolic effects of ω1.

It should be noted that despite similar RNase activity when compared to native ω1 (20), the recombinant ω1 produced by HEK-293 cells and the glycol-engineered molecules from *N. benthamiana* plants harbor significantly different N-glycosylation patterns (17), which may partly explain the different outcomes between studies. Interestingly, as compared to pWT-ω1, we observed a trend for a stronger effect on insulin sensitivity and food intake with pLe^X^-ω1, of which the glycans resemble the ones of native helminth ω1 the most.

In our study, both ω1 glycovariants were found to induce a type 2 immune response in WAT, characterized by a significant increase in Th2 cells, eosinophils and AAMs. In addition, as previously described for SEA (13), we showed that the ω1-induced increase in type 2 cytokines are clearly derived from CD4^+^ T cells, suggesting that DC-mediated Th2 skewing is required, rather than ILC2 activation, to induce WAT eosinophilia and AAM polarization. Of note, it was previously shown that pLe^X^-ω1, compared to pWT-ω1, induced a stronger Th2 polarization *in vivo* using a footpad immunization model in mice (20). In our conditions, both glycovariants induced a similar increase in the percentage of Th2 cells in metabolic tissues from obese mice, whereas pLe^X^–ω1 increased total CD4^+^ T cells to a greater extent in the liver and to a lesser extent in WAT when compared to pWT-ω1. Altogether, this suggests that the different glycans on ω1 glycovariants might lead to tissue-specific targeting of ω1 and resulting differences in total Th2 cells.

While the type 2 immune response seems not to be significantly involved in the beneficial metabolic effects of ω1, we found that treatment with both ω1 glycovariants reduced food intake, with a trend for pLe^X^-ω1 being more potent than pWT-ω1. This anorexigenic effect, which was not observed previously when mice were chronically infected with *S. mansoni* or treated with SEA (13), was dose-dependent and also observed in *Stat6*-deficient mice. Importantly, since both locomotor activity and lean body mass were not affected by ω1, this inhibition of food intake could not be due to illness induced by the treatment. Using fast-refeeding and paired feeding experiments, we clearly showed that ω1 rapidly inhibited food intake, an effect that mainly contributed to the improvements in metabolic homeostasis. Of note, in the study from Hams *et al*. using HEK-produced recombinant ω1 in the same concentration range as us, the effect of ω1 on feeding behavior and its putative contribution to the observed decrease in body weight and improvement of glucose tolerance in obese mice have not been specifically investigated (18).

Anorexia is one of the clinical manifestation of infection with different helminth species in both animals and humans. As such, deworming children infected with the hookworms *Ascaris lumbricoides* and/or *Trichuris trichuria* has been reported to increase appetite (36), suggesting a relationship between helminth infection and food intake. In rodents, infection with *Taenia taeniaformis* and *N. brasiliensis* both induced anorexia by modulating neuropeptide expression in the hypothalamus (37, 38), indicating that helminths and/or helminth products may regulate feeding behavior. The mechanism by which ω1 inhibits food intake is however still unknown and will require further neuroscience-driven approaches to be elucidated. Regulation of food intake by the central nervous system is a complex process involving both local and peripheral neuro-immuno-endocrine inputs that are mainly integrated in the hypothalamic arcuate nucleus and the brain stem *nucleus tractus solitarius* (32, 33). In our study, we did not detect accumulation of radioactively-labelled ω1 in the brain 24 hours after intraperitoneal injection, suggesting that the glycoprotein may exerts its anorexigenic effects via peripheral rather than central action(s). Upon meal ingestion, several anorexigenic peptides and hormones are produced by metabolic organs, including adipose tissues and the intestine, and can either directly act on specific neurons after crossing the blood-brain barrier or signal from the periphery via vagal nerve-mediated pathways that contribute to satiety regulation (39, 40). Leptin is a key adipose tissue-derived anorexigenic hormone which signals through the leptin receptor in the arcuate nucleus of the hypothalamus to reduce food intake (32). During obesity, hypothalamic inflammation triggers leptin resistance, resulting in increased energy consumption in obese mice (41-43). However, despite some evidence of improved systemic leptin sensitivity by ω1 (*data not shown*), we found that its anorexigenic and metabolic effects were still present in leptin receptor-deficient mice, allowing us to exclude a significant contribution of peripheral/central leptin signaling. Among the peripheral signals that regulate feeding behavior, it would be interesting to explore the involvement of a gut-brain axis, notably through vagal nerve ablation (40). Recently, *N. brasiliensis* infection and its products were also shown to increase production of the neuropeptide Neuromedin U by mucosal neurons, allowing the host to mount an effective type 2 immune response (44-46). Neuromedin U also has anorexigenic effects (47), thus it is tempting to speculate that some helminth molecules may indirectly trigger anorexia through neuro-immune interactions in the gut.

It is worth mentioning that ω1 also increased IL-13 producing Th2 cells in the liver, but, unlike SEA (13), promoted CD11c expression in Kupffer cells while not affecting the expression of YM1, suggesting that macrophages are rather polarized towards a pro-inflammatory state in this tissue. An increase in hepatic expression of fibrotic gene markers and circulating ALAT levels was also observed, both indicating increased liver damage induced by ω1. Interestingly, the ω1-induced increase in IL-13^+^ Th2 cells and IL-13 gene expression in the liver were markedly reduced in *Stat6*-deficient mice, which was accompanied by a decreased expression of fibrotic gene markers. Collectively, these findings confirmed previous studies describing that IL-13 plays a role in the development of liver fibrosis (30, 31), and that ω1 has cytotoxic effects in the liver (28, 29).

In conclusion, we report here that the helminth glycoprotein ω1 improved metabolic homeostasis in insulin-resistant obese mice by a mechanism not dependent of its type 2 immunity-inducing capacity, but rather mostly attributable to leptin receptor-independent inhibition of food intake. Further studies are required to unravel such underlying mechanisms, notably exploring the role of gut hormones on peripheral and/or central regulation of feeding behavior. Of note, with regards to its putative therapeutic potential for metabolic disorders, it is important to underline that despite beneficial effects on whole-body metabolic homeostasis, ω1 also induced early markers of mild hepatic fibrosis, partly through a type 2 immunity-mediated mechanism. Finally, by contrast to ω1, the complex mixture of SEA does not have detrimental effects on the liver and improves metabolic homeostasis through a STAT6-mediated type 2 immune response, suggesting that it may contain some other unidentified molecules, such as Dectin 2 ligands (48), with potentially beneficial immunometabolic properties.

## Acknowledgements

The authors thank Gerard van der Zon and Tessa Buckle (Leiden University Medical Center, Leiden, the Netherlands), and Uma Mahesh Gundra, Ada Weinstock, Jian-Da Lin and Mei San Tang (New York University School of Medicine, New York, USA) for their invaluable technical assistance. The authors also thank Ko Willems van Dijk and Patrick Rensen (Leiden University Medical Center) for allowing the use of the LUMC metabolic phenotyping platform (MRI and metabolic cages).

## List of abbreviations

AAM: Alternatively activated macrophage
ALAT: Alanine aminotransferase
BAT: Brown adipose tissue
DC: Dendritic cell
EE: Energy expenditure
GTT: Glucose tolerance test
HEK: Human embryonic kidney
HFD: High-fat diet
HOMA-IR: Homeostatic model assessment of insulin resistance
IFN-γ: Interferon gamma
IL: Interleukin
ILC: Innate lymphoid cell
ITT: Insulin tolerance test
LBM: Lean body mass
LFD: Low-fat diet
NAFLD: Non-alcoholic fatty liver disease
NASH: Non-alcoholic steatohepatitis
pLe^X^-ω1: Recombinant omega-1 from Le^X^-glyco-engineered *Nicotiana benthamiana* plants
PMA: Phorbol myristate acetate
pWT-ω1: Recombinant omega-1 from wild-type *N. benthamiana* plants
SEA: *Schistosoma mansoni* soluble egg antigens
STAT: Signal transducer and activator of transcription
SVF: Stromal vascular fraction
Th: T helper
TNF: Tumor necrosis factor
UCP1: Uncoupling protein 1
WAT: White adipose tissue
WT: Wild-type

